# Natural diversity of the malaria vector *Anopheles gambiae*

**DOI:** 10.1101/096289

**Authors:** The Anopheles gambiae 1000 Genomes Consortium

## Abstract

The sustainability of malaria control in Africa is threatened by rising levels of insecticide resistance, and new tools to prevent malaria transmission are urgently needed. To gain a better understanding of the mosquito populations that transmit malaria, we sequenced the genomes of 765 wild specimens of *Anopheles gambiae* and *Anopheles coluzzii* sampled from 15 locations across Africa. The data reveal high levels of genetic diversity, with over 50 million single nucleotide polymorphisms across the 230 Mbp genome. We observe complex patterns of population structure and marked variations in local population size, some of which may be due at least in part to malaria control interventions. Insecticide resistance genes show strong signatures of recent selection associated with multiple independent mutations spreading over large geographical distances and between species. The genetic variability of natural populations substantially reduces the target space for novel gene-drive strategies for mosquito control. This large dataset provides a foundation for tracking the emergence and spread of insecticide resistance and developing new vector control tools.

---

Blood-sucking mosquitoes of the Anopheles gambiae species complex exert a heavy toll on human health, being the principal vectors of *Plasmodium falciparum* malaria in Africa. Increased use of insecticide-treated bed nets (ITNs) and other methods of vector control have led to substantial reductions in the burden of malaria in Africa over the past 15 years^1,2^. However, these gains could be reversed by insecticide resistance that is rapidly spreading across the continent^3,4^ and by behavioural adaptations which cause mosquitoes to avoid contact with insecticides^5^. New insecticides are being developed for use in public health^6,7^ and there is growing support for gene drive technologies for malaria vector control^8–10^. However, relatively little is known about natural genetic diversity of Anopheles vector species, or the evolutionary and demographic processes that allow adaptive mutations to emerge and spread through mosquito populations. This knowledge is needed to maximize the efficacy and active lifespan of new insecticides and to design gene drive systems that work in the field. The Anopheles gambiae 1000 Genomes Project^†^ (Ag1000G) was established to discover natural genetic variation within this species complex, and to provide a fundamental resource for applied research into malaria vector control. Here we report on the first phase of the project, that has generated genome-wide data on nucleotide variation in 765 wild-caught mosquitoes, sampled from 15 locations in 8 countries spanning a variety of ecological settings, including rainforest, inland savanna and coastal biomes (Supplementary Fig. 1). We sampled the two major malaria vector species within the species complex, *Anopheles gambiae sensu stricto* and *Anopheles coluzzii*, which are morphologically indistinguishable and often sympatric but may differ in geographical range^11^, larval ecology^12^, behaviour^13^ and strategies for surviving the dry season^14^. *An. gambiae* and *An. coluzzii* have been classified as different species^15^ because they are genetically distinct^16–18^. However, although they undergo assortative mating^19^, reproductive isolation is incomplete: hybrids are viable and fertile, and there is evidence for hybridization in nature varying over space^20–22^ and time^23^, creating opportunities for gene flow between species^24,25^. The diversity of sampling in this project phase over geography, ecology and species is not exhaustive, but does provide a broad platform from which to explore the factors shaping mosquito population variation, evolution and speciation.

## Genomic variation

We used the Illumina HiSeq platform to perform whole genome deep sequencing on individual mosquitoes. After removing samples with low coverage (<14X) we analyzed data on 765 wild-caught specimens and a further 80 specimens comprising parents and progeny from 4 lab crosses (Supplementary Fig. 1). Sequence reads were aligned against the AgamP3 reference genome^27^ and putative single nucleotide polymorphisms (SNPs) were called from the alignments^28,29^ (Supplementary Text). The alignments were also used to identify genome regions accessible to SNP calling, where short reads could be uniquely mapped and there was minimal evidence for structural variation^30,31^. We classified 61% (141 Mbp) of the AgamP3 chromosomal reference sequence as accessible, including 91% (18 Mbp) of coding and 59% (123 Mbp) of non-coding positions (Supplementary Fig. 2A). Mendelian errors in the crosses were used to guide the design of filters to remove poor quality variant calls. In total 52,525,957 SNPs passed all quality filters. We then used statistical phasing, combined with information from sequence reads^32^, to estimate haplotypes for all wild-caught individuals. To assess the reliability of this dataset, we performed capillary sequencing of 5 genes, from which we estimated a false discovery rate of less than 1% and a sensitivity of 94% to detect SNPs within the accessible genome. We also obtained >98% concordance of heterozygous genotype calls in comparisons with capillary sequence data and >97% concordance in a second validation experiment using genotyping by primer-extension mass spectrometry^33^. We assessed phasing performance for wild-caught individuals by comparison with haplotypes generated from the crosses (Supplementary Fig. 3A) and from male X chromosome haplotypes, obtaining results comparable to human sequencing studies^32^ (Supplementary Fig. 3B).

Individual mosquitoes carried between 1.7 and 2.7 million variant alleles, with no systematic difference observed between species (Supplementary Fig. 4A). SNPs were mostly biallelic, but 21% had three or more alleles, and we discovered one variant allele every 2.2 bases of the accessible genome on average.

Variant allele density was similar on all chromosomes but markedly reduced in pericentromeric regions, as expected due to linked selection in regions of low recombination^34–36^ (Fig. 1A). Gene structure had a strong influence on nucleotide diversity, with the lowest diversity observed at non-degenerate coding positions and at the dinucleotide core of intron splice sites, as expected due to purifying selection on deleterious functional mutations (Supplementary Fig. 4B). We also found that diversity at fourfold degenerate codon positions and within short introns was twice the level found in longer introns and intergenic regions, similar to studies in *Drosophila*^37^ and *Heliconius*^38^, indicating that most non-coding sequence is under moderate selective constraint.

**Figure 1.**
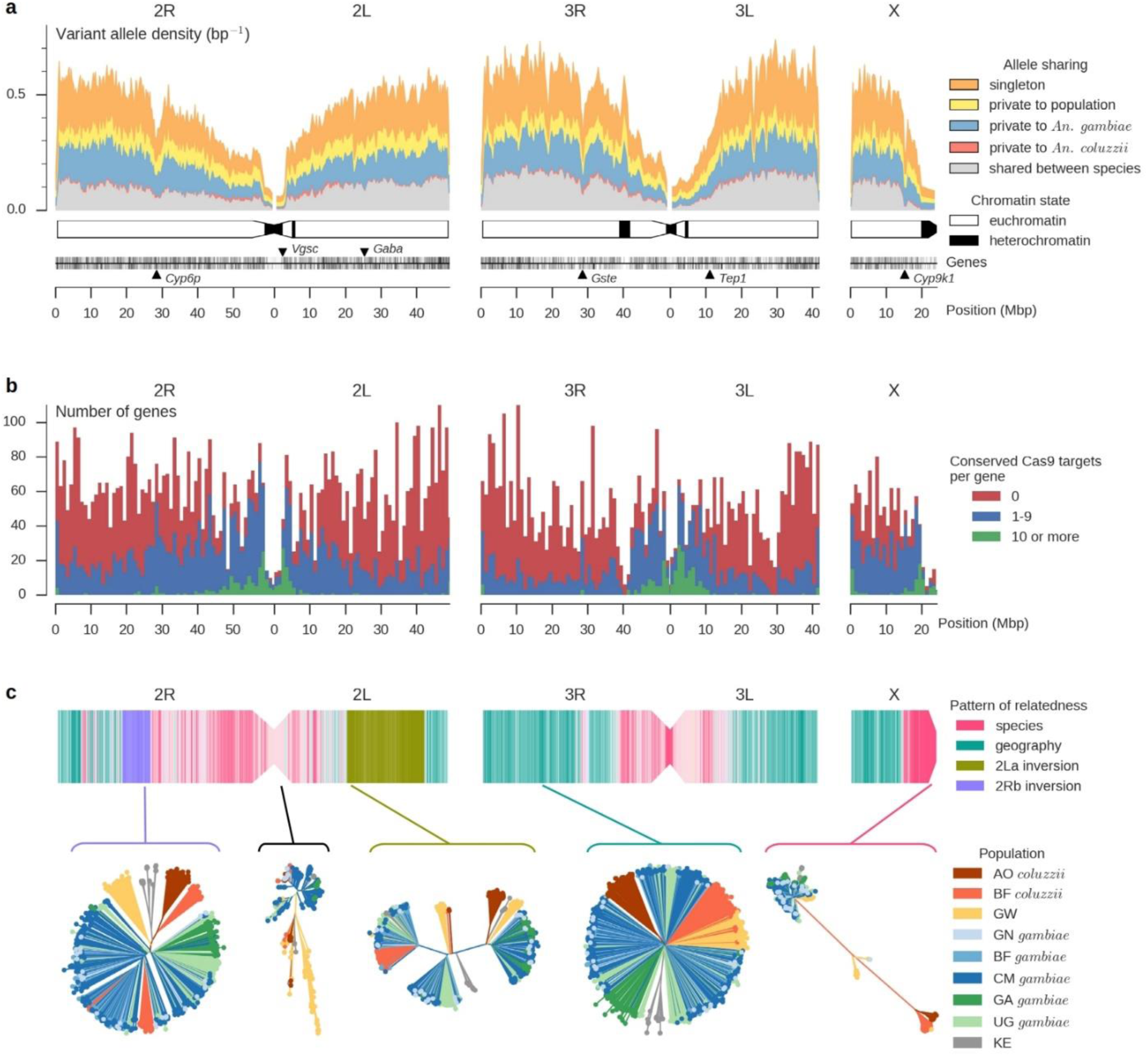
Patterns of genomic variation. **a**, Density of variant alleles in non-overlapping 200 kbp windows over the genome, computed as the number of variant alleles discovered in SNPs passing all quality filters divided by the number of accessible positions. Schematic of chromosomes below shows regions of heterochromatin^26^. **b**, Genomic distribution of genes containing conserved regions that could be targeted by CRISPR/Cas9 gene drive. **c**, Variations in the pattern of relatedness between individual mosquitoes over the genome. The upper part of the plot shows a schematic of the three chromosomes painted using colours to represent the major pattern of relatedness within non-overlapping 100 kbp windows. Below, neighbour-joining trees are shown from a selection of genomic windows that are representative of the four major patterns of relatedness found, as well as for the window spanning the Vgsc gene on chromosome arm 2L which has a unique pattern of relatedness. The strength of colour indicates the strength of the correlation with the closest major pattern. AO = Angola; BF = Burkina Faso; GW = Guinea-Bissau; GN = Guinea; CM = Cameroon; GA = Gabon; UG = Uganda; KE = Kenya. Species status is uncertain for GW and KE populations.

Since the advent of efficient genome editing using the CRISPR/Cas9 system^39^, the push to implement gene drive in *Anopheles* to carry out population suppression^10^ or replacement^9^ has intensified. However, variants within the short ~21 bp Cas9 target site represent potential resistance alleles, and thus the sheer density of SNPs could negatively impact successful deployment of gene drive in *Anopheles*. We explored the accessible coding genome for CRISPR/Cas9 target sites and found viable targets in 10,711 of 12,901 annotated genes (Supplementary Text). However, only 5,012 genes retained at least one viable target after accounting for variation within target sites, and this is likely to worsen with further population sampling (Supplementary Text). These possible target genes were spread non-uniformly across the genome, falling predominantly in pericentromeric regions, where levels of variation were lower (Fig. 1B). The evolution of resistance to gene drive will be caused both by natural variation and by the DNA repair machinery itself, and therefore drive-based methods are unlikely to work unless multiple genes and multiple sites within each gene are targeted. To that end, we identified 544 genes that each contain at least 10 non-overlapping conserved target sites, including 9 putative sterility genes^10^ (Supplementary Text). The genome sequences presented here are a valuable resource for prioritizing genes and designing gene drive strategies that will be effective in natural populations.

## Population structure and gene flow

Analysis of genetic structure provides a foundation for studying the evolutionary and demographic history of populations, and for understanding how genetic variants move between populations. We are particularly interested in gene flow across geographical ranges via migration, and gene flow between species via hybridization, as both can play a role in the spread of medically-important variants, including insecticide resistance mutations^24,25^ and introduced genetic modifications. Previous studies of the *Anopheles gambiae* complex have shown that phylogenetic relationships can vary dramatically between different genomic regions^24,25,40–42^. We therefore began by computing genetic distances between individual mosquitoes and constructing neighbour-joining (NJ) trees within non-overlapping genomic windows of 100,000 accessible bases (Fig. 1C). By analyzing the correlation between genetic distances in different genomic windows, we identified four major patterns of relatedness, systematically associated with different genomic regions. Within pericentromeric regions of chromosomes X, 3, and arm 2R, mosquitoes segregated into two distinct and widely separated clusters, largely corresponding to the two species as determined by conventional molecular diagnostics^18,43^. Individuals from coastal Guinea-Bissau, where reproductive isolation between species is believed to have broken down^20–22^, were an exception, being found in both clusters with poor correspondence to species assignments, as well as in an intermediate cluster. The large chromosomal inversions^44^ 2La and 2Rb were each associated with a distinct pattern of relatedness, as expected if gene flow is limited by reduced recombination between inversion karyotypes^42,45^. Genetic structure was weak throughout most of the remainder of the genome, with some separation of populations at the extremes of the geographical range (Angola, Kenya), but no evidence of clustering by species. In addition to these four major patterns of relatedness, we found other distinct patterns within some isolated genome regions, including windows near the voltage-gated sodium channel *(Vgsc)* gene^46^, a known locus of resistance to DDT and pyrethroid insecticides^24,25^.

These other patterns were characterized by short genetic distances between individuals from different populations and species, indicating the influence of recent selective sweeps and adaptive gene flow.

To investigate the influence of geography on population structure, we analyzed data from Chromosome 3, which is free from high frequency polymorphic inversions^44^ (Fig. 2A). We used ADMIXTURE to model each individual as a mixture deriving from *K* ancestral populations^47^ and compared with results from principal components analysis (PCA) and allele frequency differentiation (*Fst*) (Supplementary Text; Fig. 2B, 2C; Supplementary Figs. 5, 6). All analyses supported five major ancestral populations, corresponding to: (i) Guinea, Burkina Faso, Cameroon and Uganda *An. gambiae*; (ii) Gabon *An. gambiae;* (iii) Kenya; (iv) Angola *An. coluzzii*; (v) Burkina Faso *An. coluzzii* and Guinea-Bissau. These results are consistent with previous evidence that the Congo Basin tropical rainforest and the East African Rift Zone are natural barriers to gene flow^44,48–51^. Within each species, we found high *F_ST_* across these barriers, exceeding the level of differentiation between the two species at a single location (Fig. 2B; Supplementary Fig. 6B), indicating that ecological discontinuities may have a stronger impact on gene flow than assortative mating in sympatric populations.

**Figure 2.**
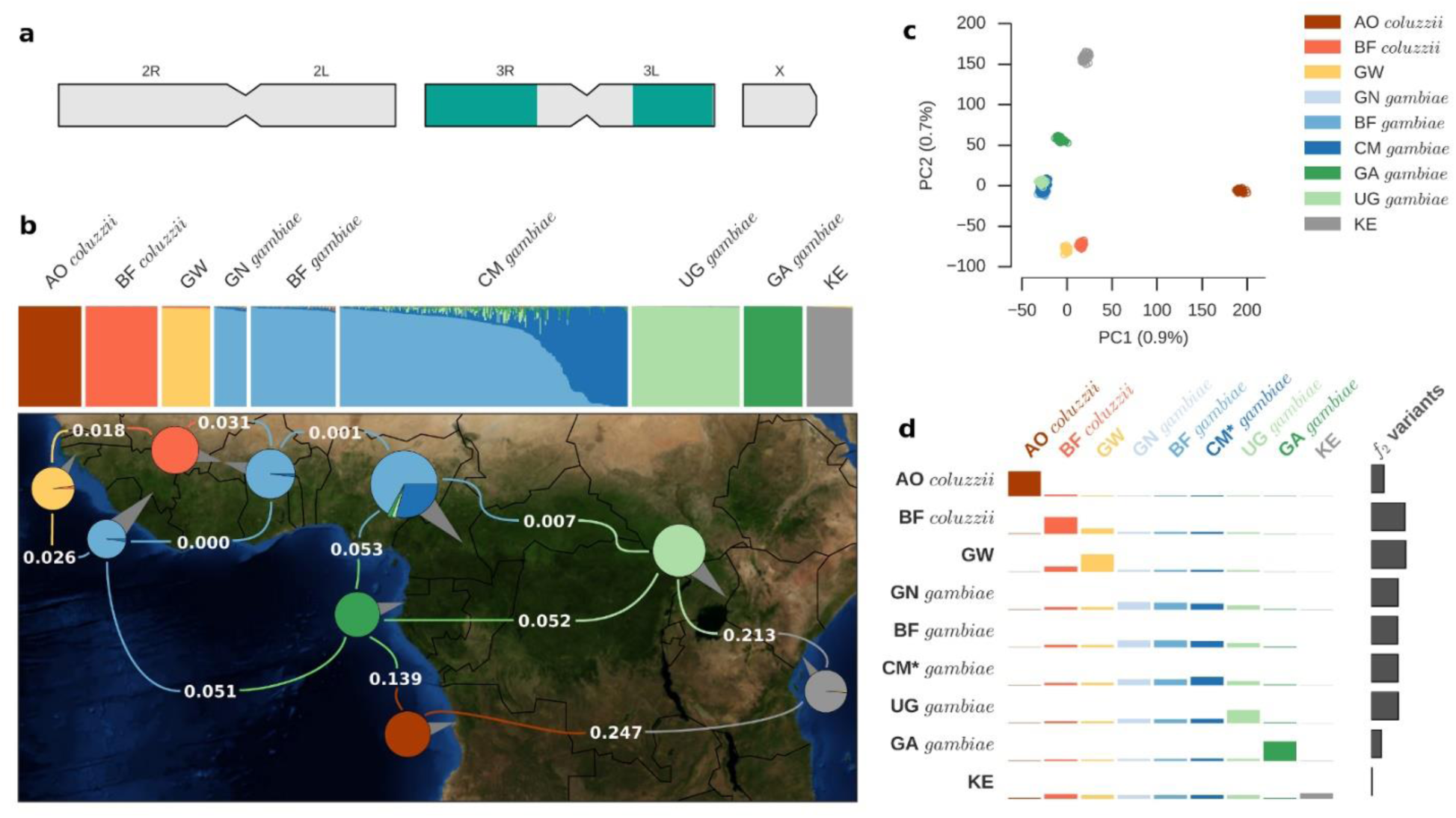
Geographical population structure. **a**, Schematic showing regions of the genome used for analyses of geographical population structure highlighted in turquoise. **b**, ADMIXTURE and allele frequency differentiation (F_ST_). The upper panel depicts each of the 765 wild-caught mosquitoes as a vertical bar, painted by the proportion of the genome inherited from each of K=8 inferred ancestral populations. Pie charts on the map depict the same inferred ancestral proportions summed over all individuals for each of 9 groups defined by species and country of origin; grey pointers attached to each pie chart show the sampling location. Average F_ST_ values are overlaid in white for selected pairs of populations. **c**, Principal components analysis. Each marker represents an individual mosquito, projected onto the first two principal components of genetic variation. **d**, Allele sharing in f_2_ variants. The height of the coloured bars represent the probability of sharing a doubleton allele between two populations. Heights are normalized row-wise for each population. Grey bars at the end of each row depict the total number of doubletons found in individuals from the given population. CM* = Cameroon savanna sampling site only.

The movement of mosquitoes affects not only the spread of genetic variants in vector populations, but also the spatial and temporal dynamics of malaria parasite transmission. Previous studies have suggested that purposeful movement of individual *Anopheles* mosquitoes is limited to short-range dispersal up to 5km^52,53^; however, recent studies have provided evidence of long-distance seasonal migration in *An. gambiae*^14^. If mosquitoes only travel short distances, we would expect to observe some differentiation between mosquitoes sampled from different geographical locations. To complement ADMIXTURE, PCA and *F_ST_* results, we also studied the sharing of rare alleles (Fig. 2D), which should be enriched for recent mutations and thus provide high resolution to detect subtle population structure. All analyses provided evidence for differentiation between Uganda and *An. gambiae* populations to the west, and between Guinea-Bissau and *An. coluzzii* from Burkina Faso (Fig. 2D; Supplementary Figs. 5, 6). However, we found no evidence for differentiation between *An. gambiae* from Guinea and Burkina Faso by any method. Some differentiation was detectable between *An. gambiae* from Burkina Faso and Cameroon, but mosquitoes were sampled from multiple sites within Cameroon along an ecological cline from savanna into rainforest, and there was evidence for some population structure and admixture associated with these different ecosystems (Fig 2B; Supplementary Figs. 5A, 6A). Considering only the Cameroon savanna site, differentiation between Cameroon and *An. gambiae* populations to the west was extremely weak (Fig 2D; Supplementary Fig. 6B). These findings are consistent with substantial rates of long-distance movement between savanna *An. gambiae* populations in West and Central Africa.

To examine gene flow between species in more detail, we analyzed a set of 506 SNPs previously found to be highly differentiated between the two species in Mali^18^. These ancestry-informative markers (AIMs) showed that a block of *An. gambiae* ancestry towards the centromere of chromosome arm 2L has introgressed into *An. coluzzii* populations in both Burkina Faso and Angola (Supplementary Fig. 7). This genomic region spans the *Vgsc* gene, where introgression of resistance mutations has previously been reported in Ghana^24^ and Mali^25^, but this is the first evidence that introgressed mutations have spread to *An. coluzzii* populations south of the Congo Basin rainforest. AIMs also showed that all mosquitoes from Guinea-Bissau carried a mixture of *An. gambiae* and *An. coluzzii* alleles on all chromosomes. These individuals were sampled from the coast, within a region of Far-West Africa that is believed to be a zone of secondary contact between the two species, because mosquitoes have frequently been found with a hybrid genotype at the species-diagnostic marker on the X chromosome, and other genetic data have suggested extensive introgression^20,22,54–56^. Our AIM results are consistent with this interpretation; however, PCA and ADMIXTURE analyses of chromosome 3 showed no evidence of recent admixture in Guinea-Bissau, rather grouping all individuals together in a single population separate from other West African populations of either species (Supplementary Figs. 5A, 6A). These results suggest a distinct demographic history for this population, and caution against the use of any single marker to infer species ancestry or recent hybridization. This point is reinforced by the observation that all mosquitoes sampled from coastal Kenya also carried a mixture of species alleles at AIMs on all chromosome arms, except for a 4 Mbp region of chromosome X spanning the location of the conventional diagnostic marker, where only *An. gambiae* alleles were present (Supplementary Fig. 7). This mixed ancestry was unexpected, as sympatry between *An. gambiae* and *An. coluzzii* does not extend east of the Rift Zone, where it is generally assumed that *An. gambiae, An. arabiensis* and *An. merus* are the only representatives of the *gambiae* complex^15^. There are several hypotheses that could explain our AIM results for Kenya, including recent or historical admixture with *An. coluzzii* populations,introgression with other species, or retention of ancestral variation. Further analyses and population sampling will be required to resolve these questions; however, our data clearly demonstrate that a simple *gambiae/coluzzii* species dichotomy is not sufficient to capture the rich diversity and complex histories of contemporary populations.

## Variations in population size

Demographic events in the history of a population, including expansions or contractions in effective population size (*N_e_*), can be inferred from the genomes of extant individuals^57^. For malaria vectors, inferring changes in *N_e_* has practical relevance, because it could provide a means to evaluate the impact of vector control interventions. For each population, we computed summary statistics of genetic diversity that are influenced by demographic history, including nucleotide diversity (π), site frequency spectra (SFS) and decay of linkage disequilibrium (LD) (Fig. 3A). All populations north of the Congo Basin rainforest and west of the Rift Zone had characteristics of large *N_e_* and population expansion, with high diversity (π = 1.5%), an excess of rare variants (Tajima’s *D* < −1.5) and extremely rapid decay of LD (*r*^2^ < 0. 01 within < 1kbp). In Gabon and Angola, we found lower diversity, more extensive LD, and an SFS closer to the null expectation under constant population size, indicating smaller *N_e_* and different demographic histories. In Kenya, we found the lowest level of diversity (π = 0.9%), a strong deficit of rare variants (Tajima’s *D* > 2), and much longer LD (*r*^2^ > 0.01 at 10Mbp), suggesting a recent population bottleneck.

**Figure 3.**
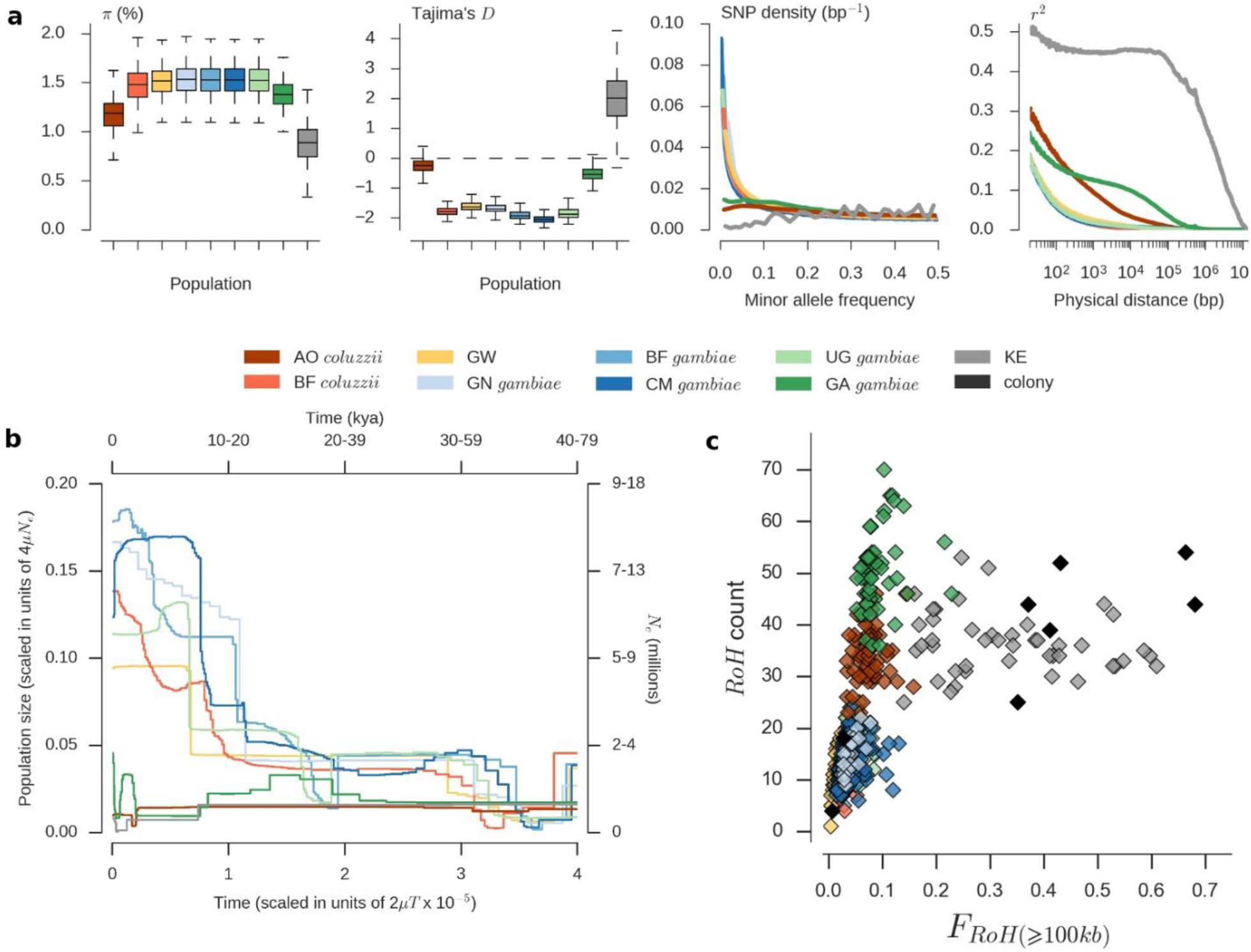
Genetic diversity and population size history. **a**, Statistics summarizing features of genetic diversity within each population. Nucleotide diversity (π) and Tajima’s D are shown as the distribution of values calculated in non-overlapping 20kbp genomic windows. SNP density depicts the distribution of allele frequencies (site frequency spectrum) for each population, scaled such that a population with constant size over time is expected to have a constant SNP density over all allele frequencies. **b**, Stairway Plot of changes in population size over time, inferred from site frequency spectra. Absolute values of time and Ne are shown on alternative axes as a range of values, assuming lower and upper limits for the mutation rate as 2.8×10^−9^ and 5.5×10^−9^ respectively, and assuming T=11 generations per year. **c**, Runs of homozygosity in individual mosquitoes, highlighting evidence for recent inbreeding in Kenyan (grey) and colony (black) mosquitoes.

We inferred the scale and timing of historical changes in *N_e_* using two methods, Stairway Plot^58^ and ∂a∂i^59^, both using site frequency spectra but taking different modelling approaches. Stairway Plot inferred a major expansion in all populations north of the Congo Basin rainforest and west of the Rift Zone (Fig. 3B; Supplementary Fig. 8A). Three-epoch ∂a∂i models also inferred expansions in these populations, with comparable magnitudes and timings (Supplementary Fig. 8B).Translating these results into absolute values for the timing and scale of expansion depends on the mutation rate, which has not been estimated in *Anopheles.* Estimates in *Drosophila*^60,61^ range from 2.8x10^−9^ to 5.5x10^−9^, which would date the onset of a major expansion in the range 7,000 to 25,000 years ago (Fig. 3B). *An. gambiae* and *An. coluzzii* are both highly anthropophilic and so should have benefited from historical human population growth, particularly the expansion of agricultural Bantu-speaking groups originating from north of the Congo Basin beginning ~5,000 years ago^62–65^. The difference in timing suggests that either the true mutation rate in *Anopheles* is higher than we have assumed, or that mosquito populations benefited from some earlier human population growth or another factor. There have also been major climatic changes since the last glacial maximum ~20,000 years ago, when overall environmental conditions in Africa were much drier than present^66^. If a general reduction in aridity was the major driver, then we might expect to see evidence for expansion in all mosquito populations sampled. However, we inferred different demographic histories in Angola, Gabon and Kenya, although more recent *N_e_* fluctuations may be obscuring earlier events in these populations, particularly in Gabon and Kenya (Fig. 3B; Supplementary Fig. 8).

In Kenya in 2006, free mass distribution of ITNs was carried out in multiple districts, resulting in a rapid increase in ITN coverage, from less than 10% in 2004 to over 60% by the beginning of 2007^67^. Mosquitoes for this study were sampled from Kilifi County in 2012, and therefore originate from populations experiencing sustained ITN pressure for several years. To investigate evidence for a very recent bottleneck in this population, we analyzed runs of homozygosity (ROH). Kenyan mosquitoes had between 10-60% of their genome within a long ROH, a level not seen in any other population (Fig. 3C). This level of homozygosity is comparable to that found in isolated human populations^68^ and domestic animal breeds^69^ due to recent inbreeding. We also observed similar ROH in mosquitoes originating from lab colonies, which are typically maintained in cages of at most a few hundred individuals, and thus where inbreeding is inevitable (Fig. 3C). Genetic signatures of recent inbreeding have previously been observed in a mosquito population from Burkina Faso^70^ and in a separate study of mosquitoes collected from Kilifi in 2010^71^. However, there remains uncertainty as to whether ITN scale-up is the root cause of mosquito population decline in Kilifi^71^, particularly as other studies have found evidence for lower *N_e_*^48^ and changes in species abundance^72^ in the region pre-dating high levels of ITN coverage. Furthermore, while ITNs have been effective in Kilifi, a substantial reduction in malaria prevalence had occurred prior to free ITN distribution^73^, thus multiple factors may be affecting vector and parasite populations in this region. Sequencing mosquitoes and parasites before, during and after interventions, and across a range of ecological and epidemiological settings, could help to resolve these questions, providing valuable information about the impact and efficacy of different control strategies.

## Evolution of insecticide resistance

Insecticide resistance is a polygenic trait with a broad phenotypic range, and several genes have previously been associated with resistance in *Anopheles,* including genes encoding insecticide binding targets and genes involved in insecticide metabolism^3^. It is not yet clear which of these genes, if any, are responsible for epidemiologically relevant levels of resistance in the field. However, mutations that confer an advantage under strong pressure from insecticide use will be positively selected, and so evidence of recent selection in natural populations can help to identify and prioritize resistance genes for further study. We used metrics of haplotype diversity^74^ (H12) and haplotype homozygosity^75^ (XP-EHH) to scan the genome for genes with evidence of recent selection. Both metrics revealed strong signals of selection in multiple populations at several genome locations containing genes associated with insecticide resistance (Fig. 4; Supplementary Fig. 9). These included *Vgsc,* confirming evidence for selection from population structure analyses described above; *Gste,* a cluster of glutathione S-transferase genes including *Gste2,* previously implicated in metabolic resistance to DDT and pyrethroids^76,77^; and *Cyp6p,* a cluster of genes encoding cytochrome P450 enzymes, including *Cyp6p3* which is upregulated in permethrin and bendiocarb resistant mosquitoes^78,79^.

**Figure 4.**
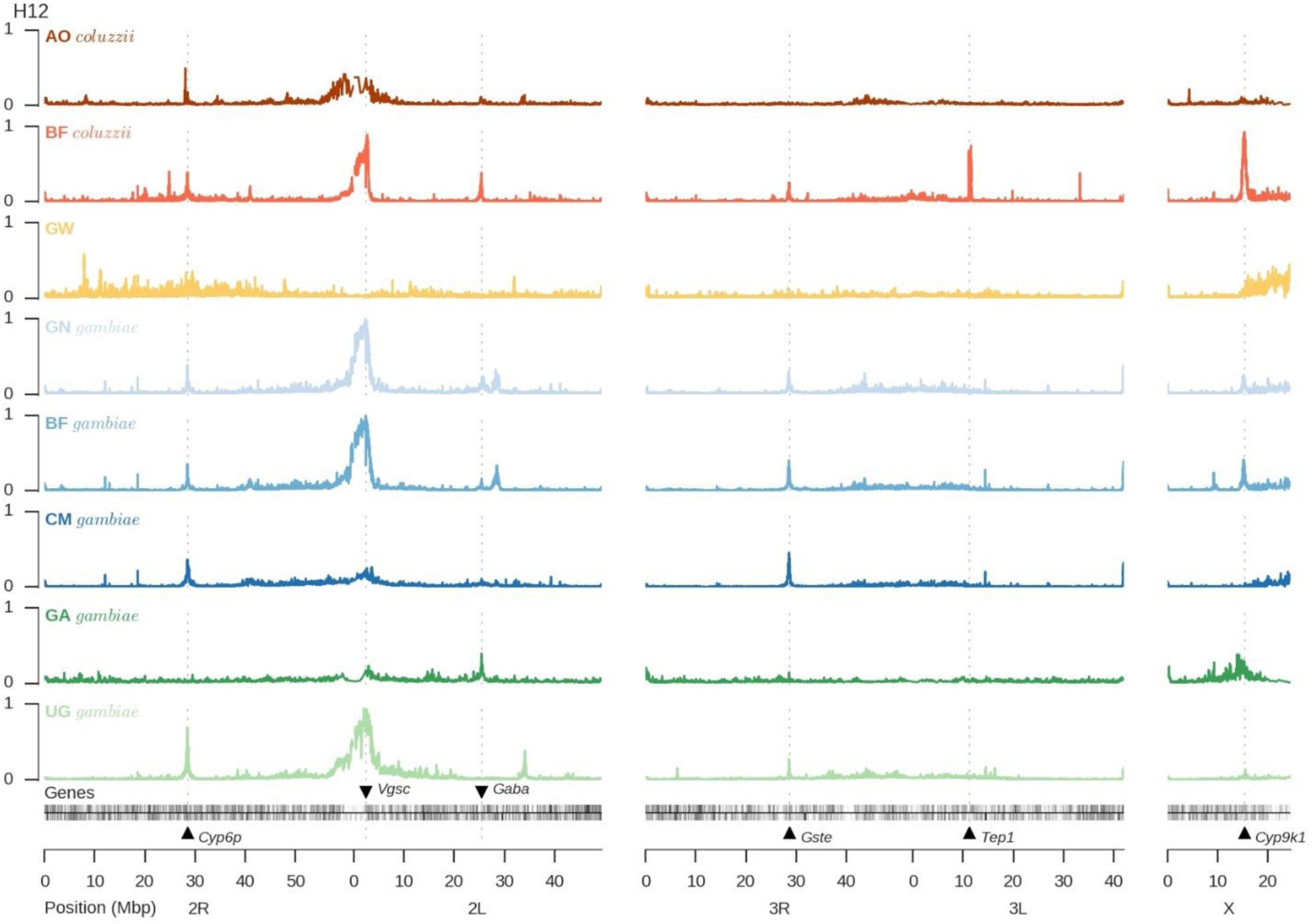
Genome scans for signatures of recent selection. Each track plots the H12 statistic in non-overlapping windows over the genome. A value of 1 indicates low haplotype diversity within a window, expected if one or two haplotypes have risen to high frequency due to recent selection. A value of 0 indicates high haplotype diversity, expected in neutral regions. Kenya is not shown because the high genome-wide levels of homozygosity mean reduced power to detect evidence of recent selection in specific genome regions.

Mutations in *An. gambiae Vgsc* codon 995 (orthologous to *Musca domestica Vgsc* codon 1014), known as *“kdr”* due to their knock-down resistance phenotype, reduce susceptibility to DDT and pyrethroids by altering binding-site conformation^46^. We found the Leucine→Phenylalanine (L995F) *kdr* mutation at high frequency in West and Central Africa (Guinea 100%; Burkina Faso 93%; Cameroon 53%; Gabon 36%;Angola 86%). A second *kdr* allele, the Leucine→Serine (L995S) mutation, was present in Central and East Africa (Cameroon 15%; Gabon 65%; Uganda 100%; Kenya 76%). To investigate the origins and movements of these two distinct *kdr* mutations, we analyzed the genetic backgrounds on which they were carried, using information from all 1,718 biallelic SNPs found across both coding and non-coding regions of the *Vgsc* gene (Fig. 5). The L995F mutation occurred in five distinct haplotype clusters (labeled F1-F5 in Fig. 5), while the L995S mutation was found in a further 5 haplotype clusters (labeled S1-S5 in Fig. 5), indicating that the number of independent origins for each of these mutations is higher than previously estimated^80–82^. Several *kdr* haplotypes have also spread between populations, despite considerable geographic distance or ecological separation. For example, haplotype F1 is present in both species and in 4 countries spanning the Congo Basin rainforest, and is the same haplotype previously found to be introgressed from *An. gambiae* into *An. coluzzii* in Ghana^24^, indicating strong selection across a variety of ecological settings. Additionally, three *kdr* haplotypes (F4, F5, S2) were found in both Cameroon and Gabon, providing multiple examples of recent adaptive gene flow between these two otherwise highly differentiated populations. Finally, the S3 haplotype was found in both Uganda and Kenya, showing that adaptive alleles can even cross the Rift Zone. While these remarkable patterns of evolution and adaptive gene flow were primarily driven by the two *kdr* mutations, we found 16 other non-synonymous mutations within *Vgsc* at a frequency above 1% (Fig. 5), of which 13 occurred exclusively on haplotypes carrying the L995F *kdr* mutation, suggesting secondary selection acting on mutations that enhance or compensate for the primary *kdr* phenotype.

**Figure 5.**
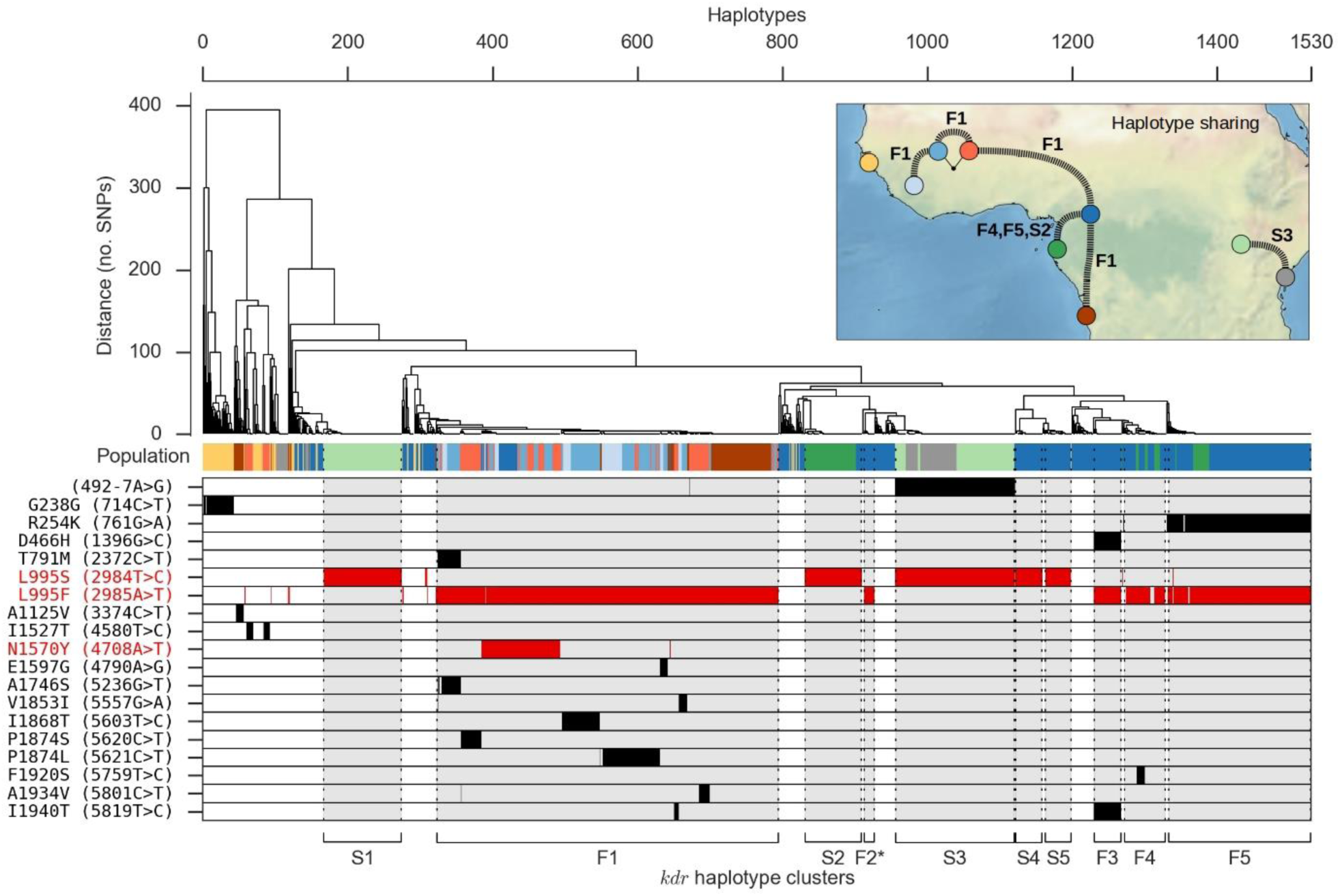
Haplotype structure at the Vgsc gene. The upper panel shows a dendrogram obtained by hierarchical clustering of haplotypes from wild-caught individuals. The colour bar immediately below shows the population of origin for each haplotype. Inset map depicts haplotypes shared between populations. The lower panel shows alleles carried by each haplotype at 19 SNPs with allele frequency > 1% that either change the amino acid sequence or occur within a splice region, and therefore may affect protein function (white = reference allele; black = alternate allele; red = previously known resistance-conferring allele). At the lower margin, we label 10 haplotype clusters carrying a kdr mutation (either L995F or L995S).

Metabolic resistance is of particular concern as it has been implicated in extreme resistance phenotypes observed in some *Anopheles* populations^79^. At both *Gste* and *Cyp6p* we found evidence that resistance has emerged multiple times, and is also spreading between species and over considerable distances. At the *Gste* locus we found at least four distinct haplotypes under selection (Supplementary Fig. 10A). One of these haplotypes carried the Gste2-I114T mutation which enhances DDT metabolism^77,83^, though the other three haplotypes did not carry any known resistance mutations. At the *Cyp6p* locus we found at least eight distinct haplotypes under selection (Supplementary Fig. 10B). Clearly there is much to learn regarding the molecular basis of metabolic resistance, and our SNP data can be used to identify candidate resistance mutations. For example, at both loci we found multiple non-synonymous SNPs that were strongly associated with haplotypes under selection (Supplementary Fig. 10). These data provide a starting point for new studies to characterize resistance phenotypes, and to develop improved tools for monitoring and responding to the emergence and spread of resistance in natural populations.

## Discussion

In this first phase of the Ag1000G project we have focused on nucleotide variation, revealing an extraordinary reservoir of natural genetic diversity in mosquito populations. Nucleotide diversity is 1.5% in most populations, twice that reported for African populations of *Drosophila melanogaster*^37,84^ and ten times greater than human populations^30^, sustained by a network of large and highly interconnected populations. The genomes that we have sequenced convey a rich mosaic of different ancestries, shaped by geography, ecology, speciation, migration, selection, recombination and chromosomal inversions, with different forces predominating in different genomic regions. Mosquito populations in different parts of Africa have experienced major demographic changes, including expansions and contractions in size, influenced at least in part by major events in the history of our own species. The introduction of insecticides has led to intense selection pressure, repeatedly driving resistance mutations to high frequency and demonstrating the potential for adaptive gene flow across the entire continent. The data we have generated provide a resource for studying and responding to the ongoing evolution of malaria vector populations. To facilitate access to this resource we have developed a novel web application^‡^ that enables visual exploration of genomic data on populations and individual mosquitoes from the scale of a whole chromosome down to individual nucleotides. Future project phases will increase both the geographical and taxonomic representation of mosquito genomes sequenced, and will explore other forms of genetic variation, including small insertion/deletion polymorphisms and large structural variation. We will also continue to study fundamental population-genetic processes, including mutation, recombination, natural selection, and the fine structure and history of gene flow between populations.

In 1899 Ronald Ross proposed that malaria could be controlled by destroying breeding sites of the mosquitoes that transmit the disease^85^. *An. gambiae,* identified in the same year by Ross as a vector of malaria in Africa^86^, has proved resilient to a century of attempts to repress it. The vector control armamentarium needs to be expanded, not only with new classes of insecticide and novel genetic control strategies, but also with more effective tools for gathering intelligence, to enable those responsible for planning and executing interventions to stay ahead of the mosquito's remarkable capacity for evolutionary adaptation. There remain major knowledge gaps, e.g., concerning the rate and range of long-distance migration, which are fundamental to understanding both malaria transmission and the spread of insecticide resistance, and which will require detailed spatiotemporal analysis of mosquito population structure. Most importantly, it is essential to start collecting population genomic data prospectively as an integral part of major vector control interventions, to identify which strategies are most likely to cause increased resistance, or what it takes to cause a population crash of the magnitude observed in our Kenyan data. By treating each major intervention as an experiment, and by analyzing its impact on mosquito populations, we can aim to improve the efficacy and sustainability of future interventions, while at the same time learning about basic processes of ecology and evolution.

## Methods

Methods are described in Supplementary Text.

## Data availability

All sequence reads from the Ag1000G project are available from the European Nucleotide Archive (ENA-http://www.ebi.ac.uk/ena) under study PRJEB1670. Submission of sequence read alignments and variant calls from Ag1000G phase 1 is in progress under ENA study PRJEB18691. Variant and haplotype calls and associated data from Ag1000G phase 1 can be explored via an interactive web application or downloaded via the MalariaGEN website (https://www.malariagen.net/projects/ag1000g#data).

## Acknowledgments

This work was supported by the Wellcome Trust (090770/Z/09/Z; 090532/Z/09/Z; 098051) and Medical Research Council UK and the Department for International Development (DFID) (MR/M006212/1). The authors would like to thank the staff of Wellcome Trust Sanger Institute Sample Logistics, Sequencing and Informatics facilities for their contributions. SO’L and AB were supported by a grant from the Foundation for the National Institutes of Health through the Vector-Based Control of Transmission: Discovery Research (VCTR) program of the Grand Challenges in Global Health initiative of the Bill & Melinda Gates Foundation.

## Supplementary figures

**Supplementary Figure 1.**
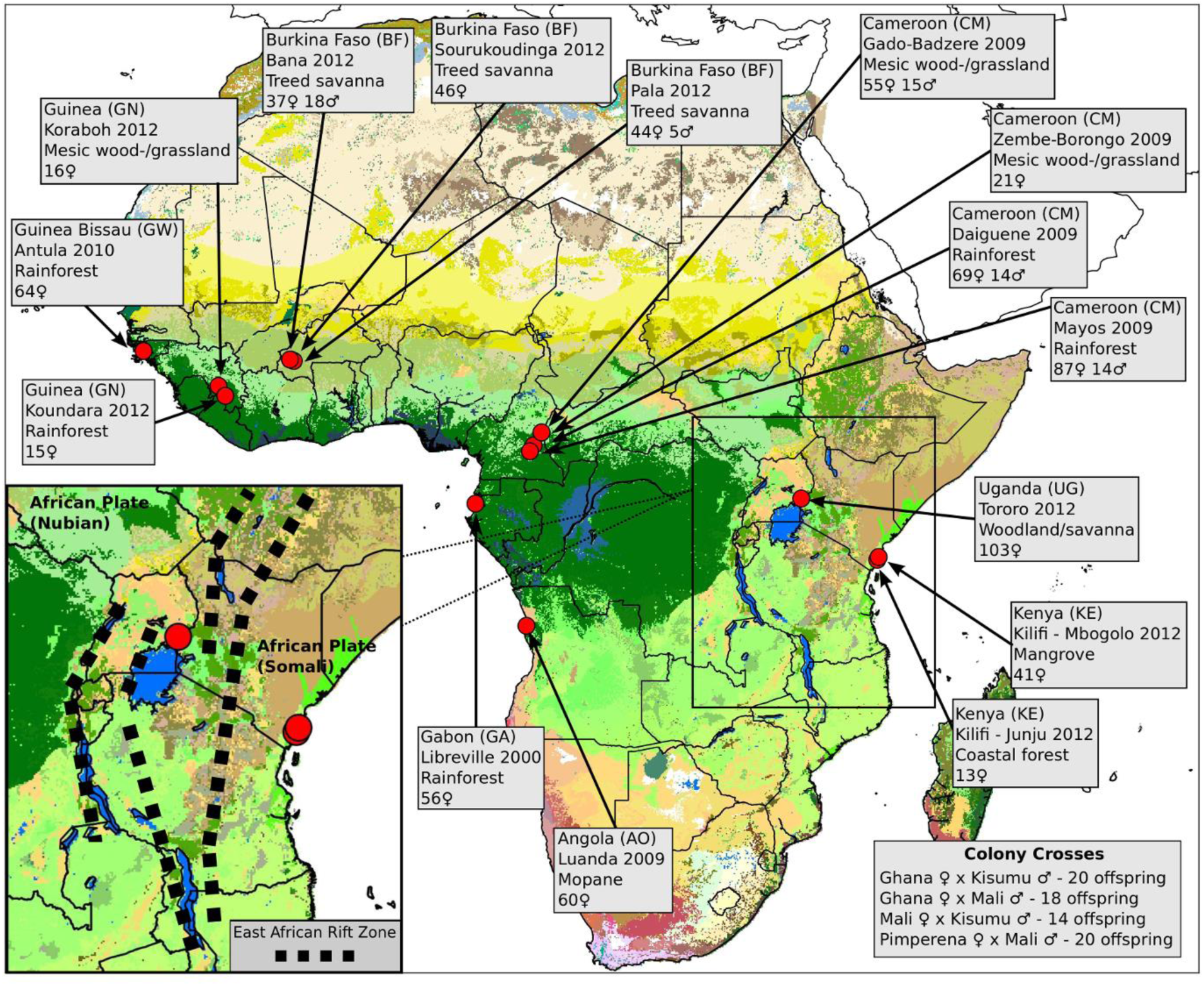
Overview of population sampling. Red circles show sampling locations for wild-caught mosquitoes; black outlines show country borders. Colours in the map represent ecosystem classes; dark green represents forest ecosystems, see (87) Fig. 9 for a complete colour legend. The Congo Basin tropical rainforest is the large region of dark green in Central Africa, spanning parts of Cameroon, Equatorial Guinea, Gabon, Central African Republic, Republic of Congo and Democratic Republic of Congo. Sampling details for each site are shown in light grey boxes, including country (two-letter country code), name of sampling site, year of collection, predominant ecosystem classification87 for the local region, and number and sex of individuals sequenced. Further details of sampling locations and methods are provided in Supplementary Text. For colony crosses, the direction of cross (colony of origin of mother and father) and number of offspring is shown. The inset map depicts geological fault lines in the East African Rift Zone^*^.

**Supplementary Figure 2.**
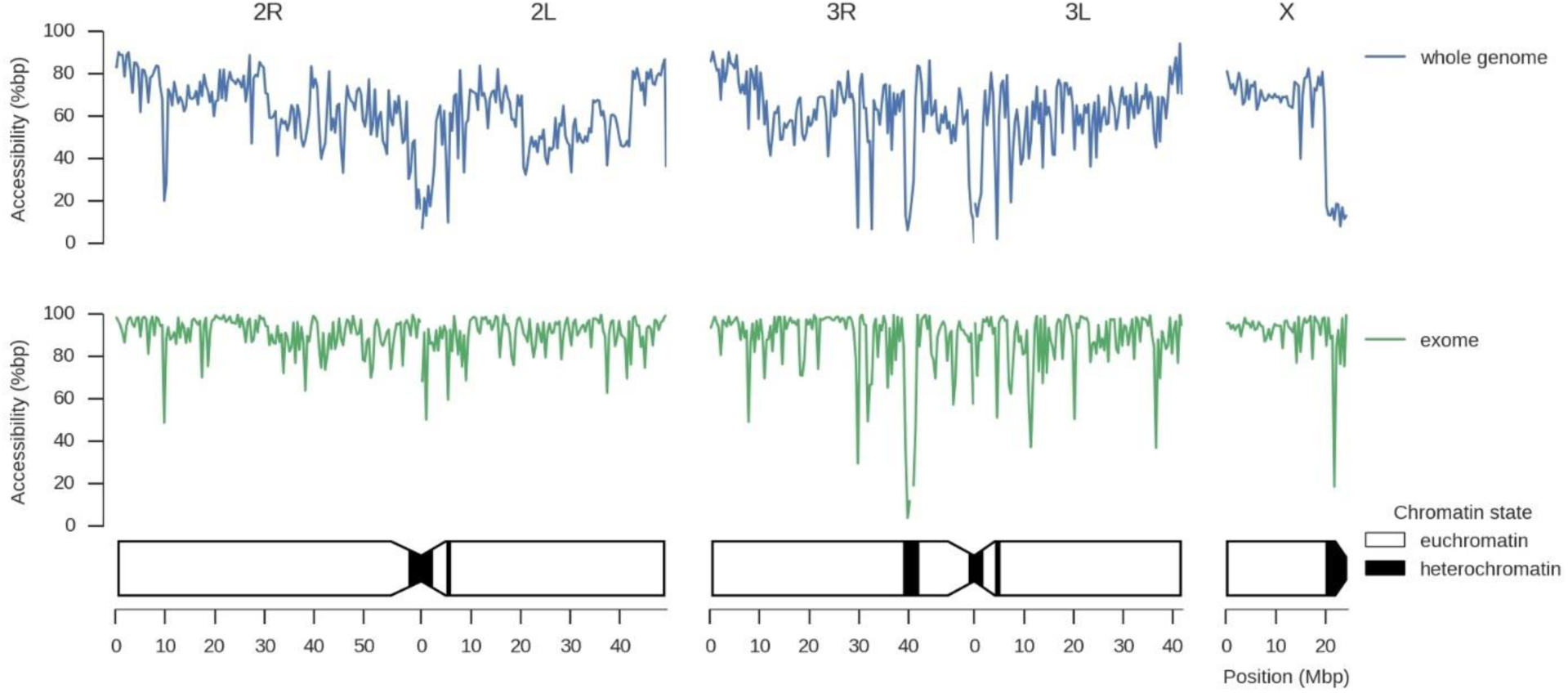
Genome accessibility. Plots show the percentage of accessible bases in non-overlapping 400kbp windows. The schematic of chromosomes below shows chromatin state predictions from (26).

**Supplementary Figure 3.**
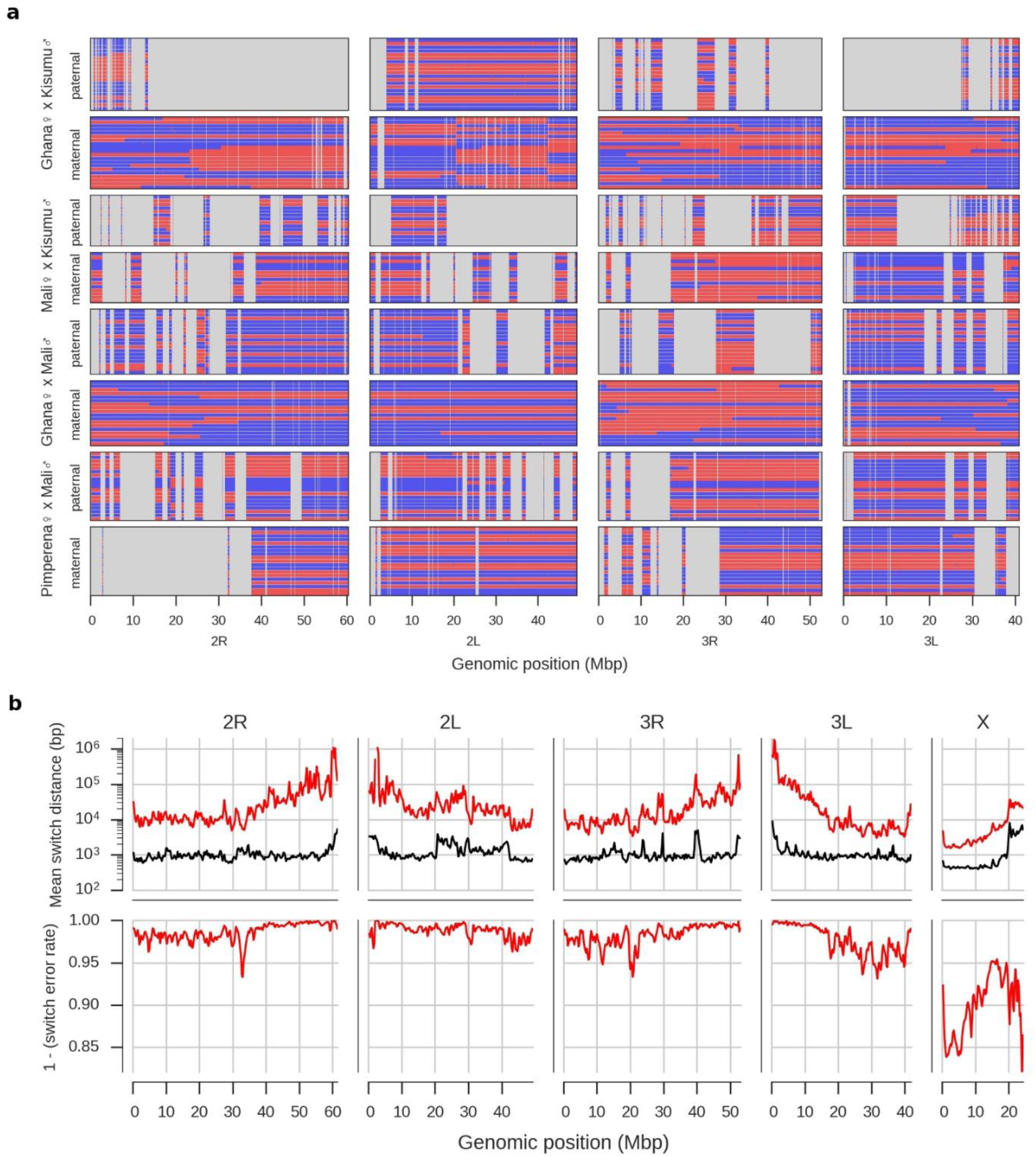
Haplotype validation. **a**, Haplotypes inferred in the crosses. Each panel shows either maternal or paternal haplotypes from a single cross. Each row within a panel represents a single progeny haplotype. Haplotypes are coloured by parental inheritance (blue = allele from parent’s first chromosome, red = allele from parent's second chromosome). Switches between colours along a haplotype indicate putative recombination events. Regions that were within a run of homozygosity in the parent and thus not informative for haplotype validation are masked in grey. **b**, Error rate estimates for haplotypes inferred in wild-caught individuals. Upper plots show estimates for the mean switch distance (red line) in windows over the genome, compared to the mean switch distance if heterozygotes were phased randomly (black line). Lower plots show the switch error rate, which estimates the probability of a switch error occurring between two adjacent heterozygous genotype calls.

**Supplementary Figure 4.**
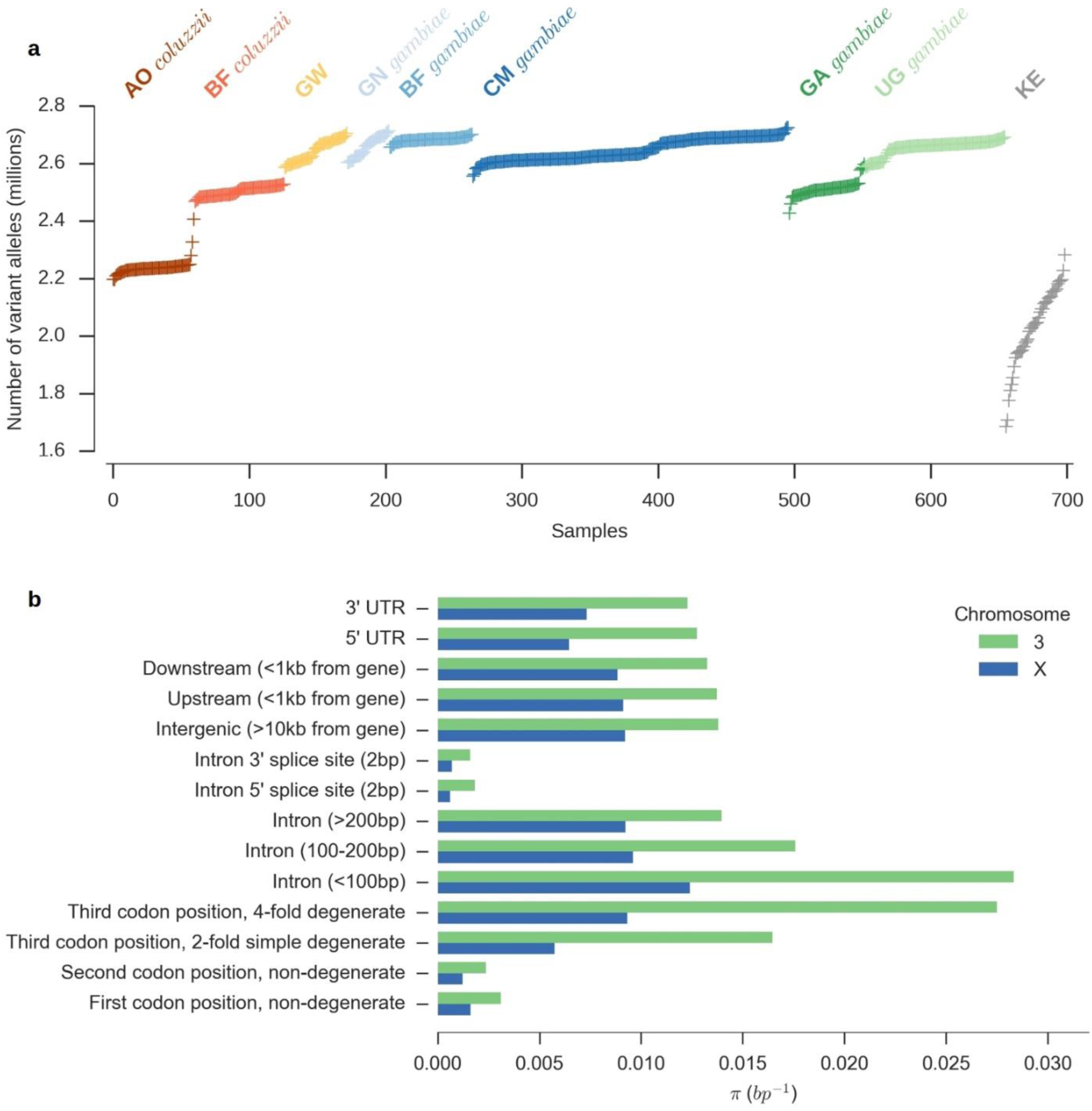
Variant discovery and nucleotide diversity. **a**, Total number of variant alleles discovered per individual mosquito sequenced. Only females are plotted. **b**, Average nucleotide diversity (π) in relation to gene architecture.

**Supplementary Figure 5.**
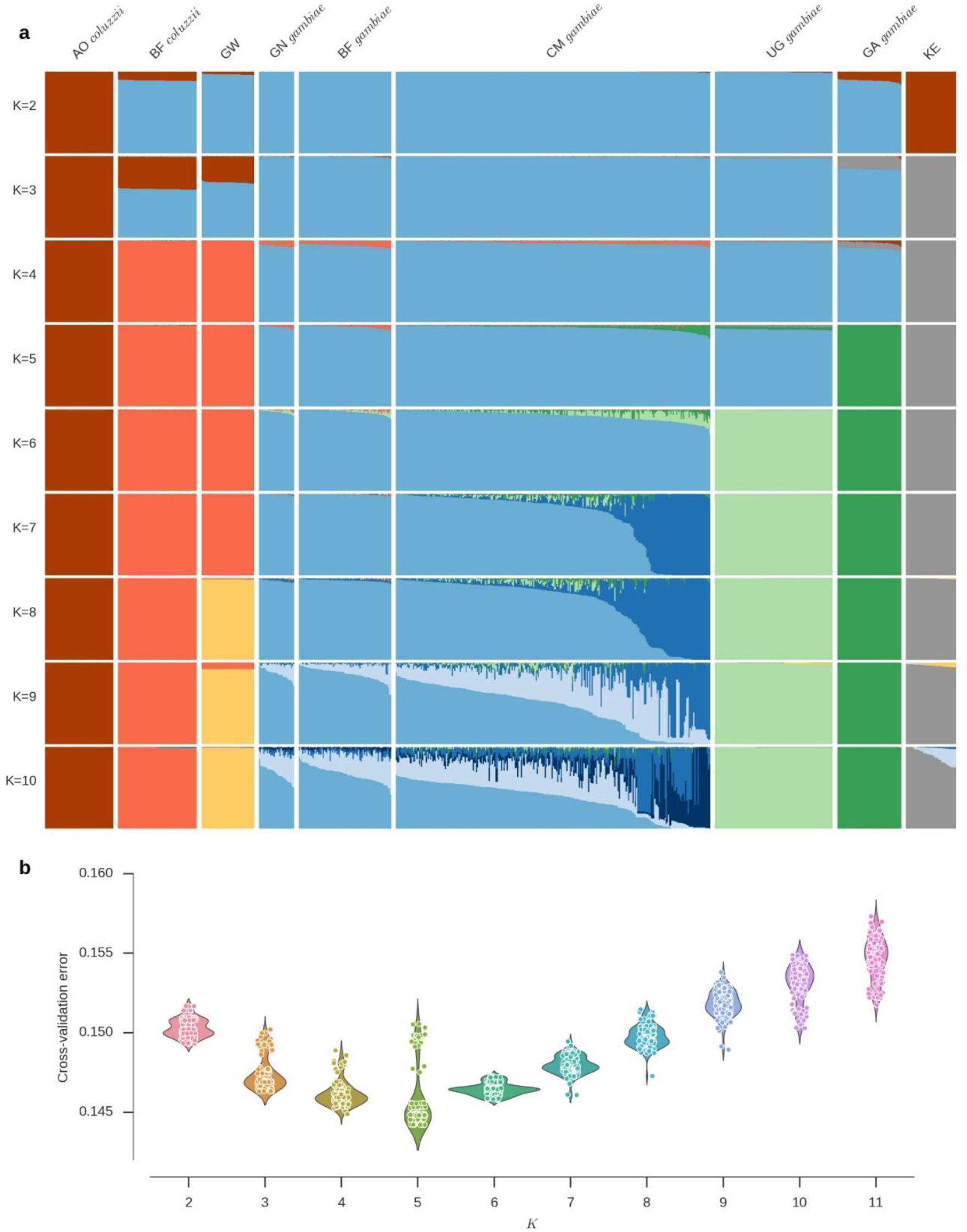
ADMIXTURE analysis. **a**, Ancestry proportions within individual mosquitoes for ADMIXTURE models from K=2 to K=10 ancestral populations. Each vertical bar represents the proportion of ancestry within a single individual, with colours corresponding to ancestral populations. These data are the average of the major q-matrix clusters derived by CLUMPAK analysis. **b**, Violin plot of cross-validation error for each of 100 replicates for each K values.

**Supplementary Figure 6.**
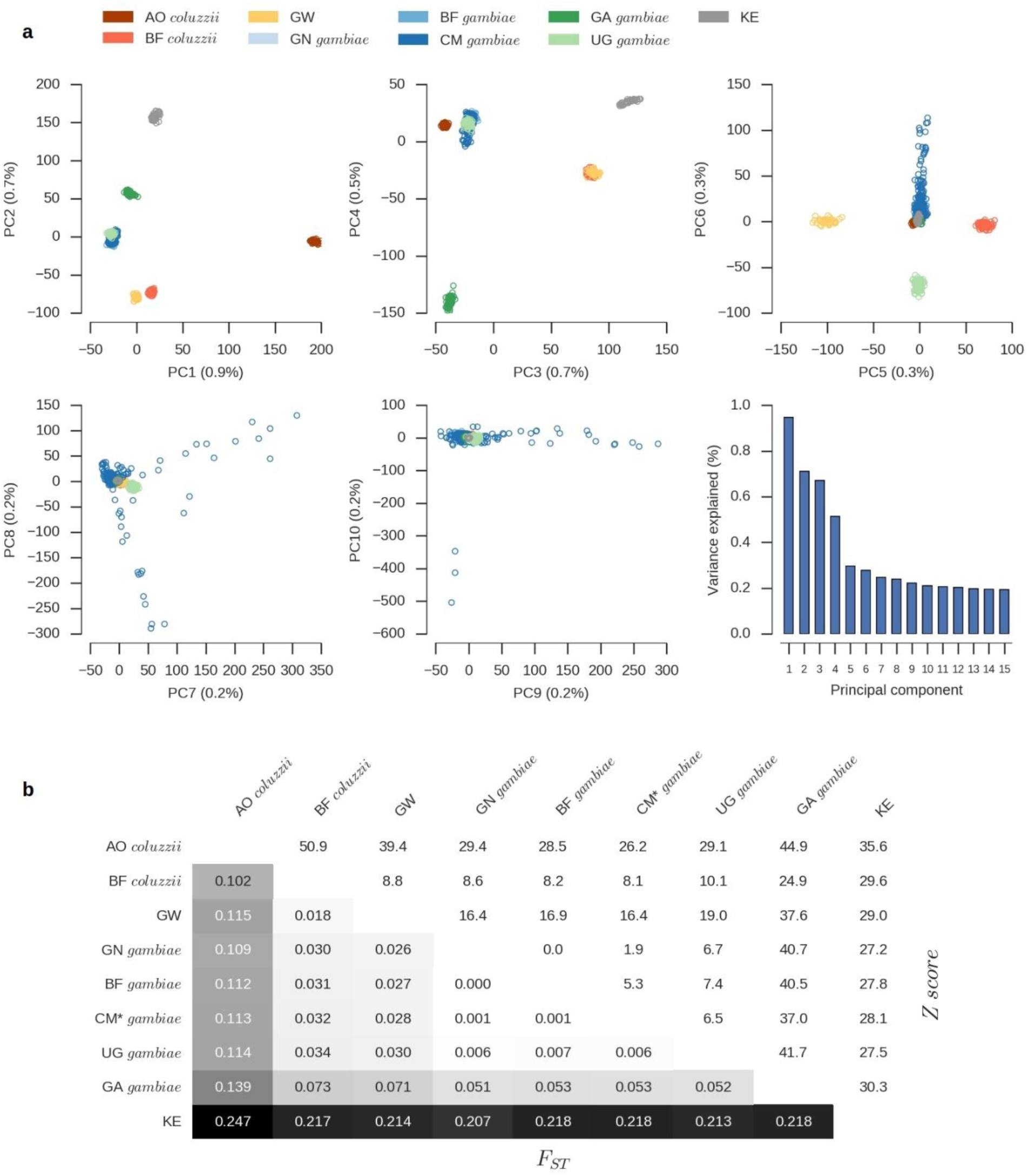
Population structure and allele frequency differentiation. **a**, Principal components analysis of the 765 wild-caught mosquitoes, showing the first 10 components of genetic variation. The final panel shows the variance explained by each component. b, Average allele frequency differentiation (F_st_) between pairs of populations. The lower left triangle shows average Fst between each population pair. The upper right triangle shows the Z score for each F_st_ value estimated via a block-jackknife procedure.

**Supplementary Figure 7.**
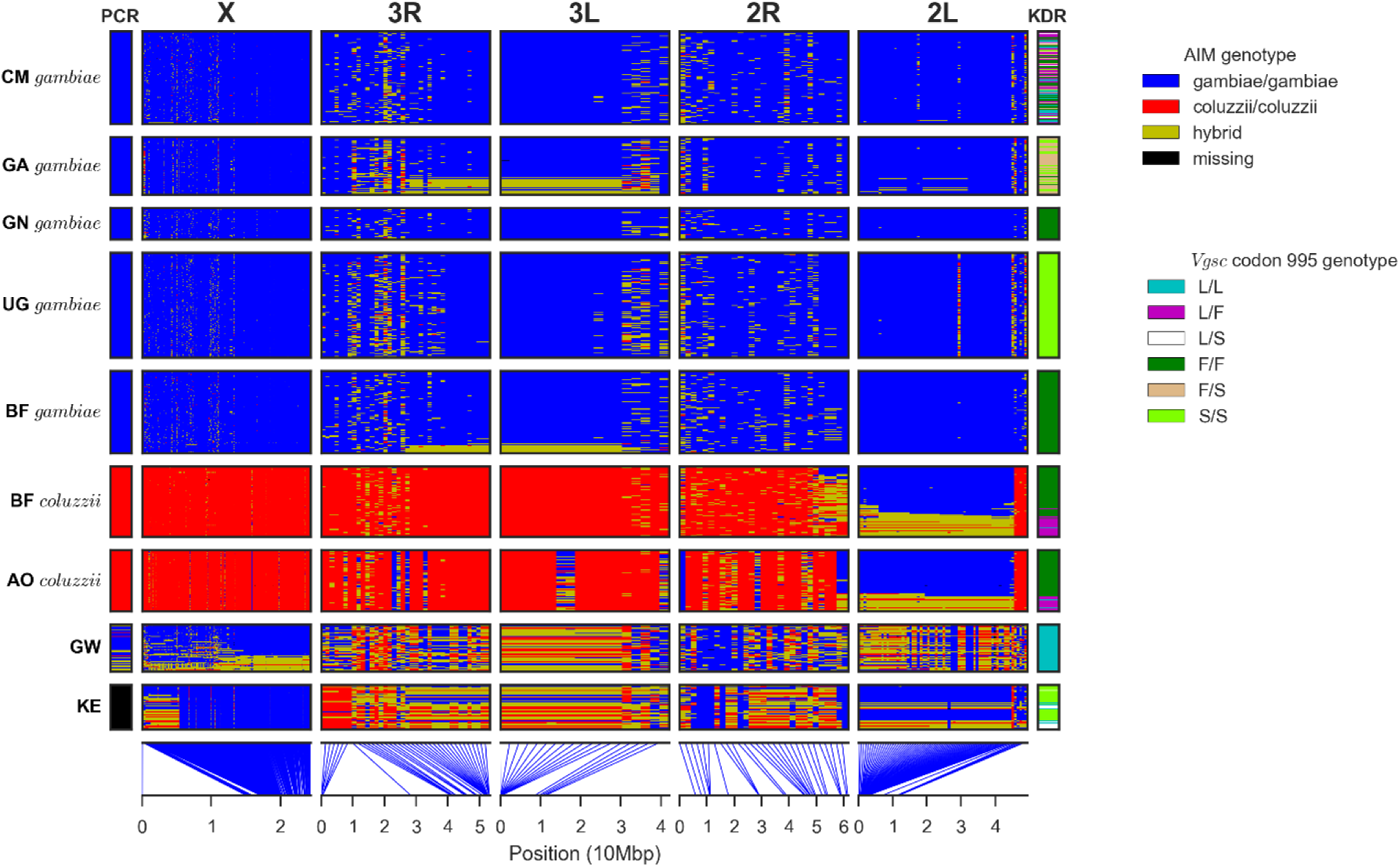
Ancestry informative markers (AIMs). Rows represent individual mosquitoes (grouped by population) and columns represent SNPs (grouped by chromosome arm). Colours represent genotype. The column at the far left shows the species assignment according to the conventional molecular test based on a single marker on the X chromosome, which was performed for all individuals except Kenya (KE). The column at the far right shows the genotype for kdr mutations in Vgsc codon 995. Lines at the lower edge show the physical locations of the AIM SNPs.

**Supplementary Figure 8.**
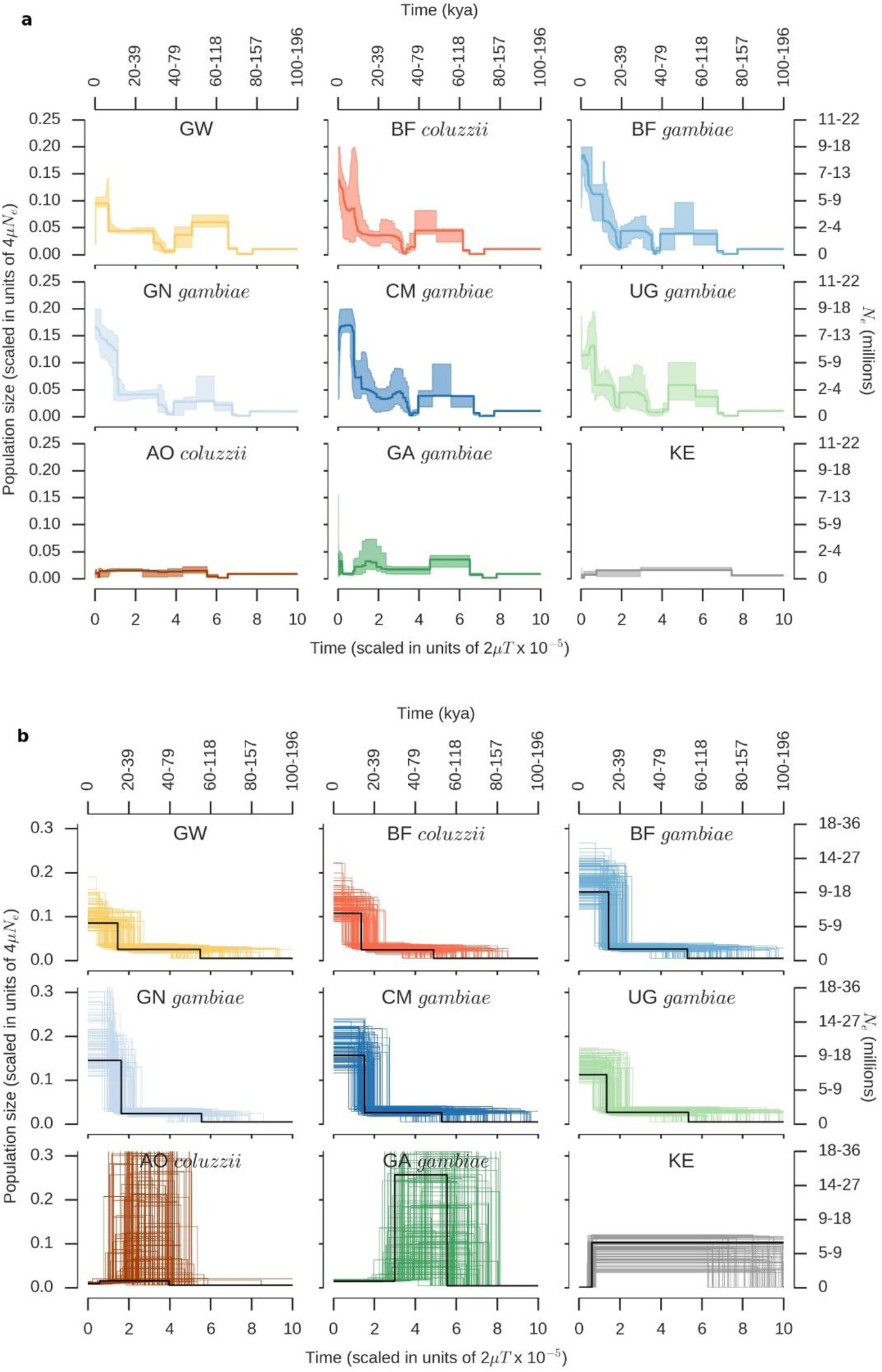
Inferred population size histories. **a**, Stairway Plot inferred histories for each population. The shaded area shows the 95% confidence interval from 199 bootstrap replicates. **b**, Inferred histories from ∂a∂i three epoch models. Black line shows the history with the highest likelihood found by optimization; coloured lines show 100 histories with the highest likelihoods from even sampling of the model parameter space (Supplementary Text). Absolute time and Ne are shown as a range assuming 11 generations per year and a mutation rate of between 2.8×10^−9^ and 5.5×10^−9^.

**Supplementary Figure 9.**
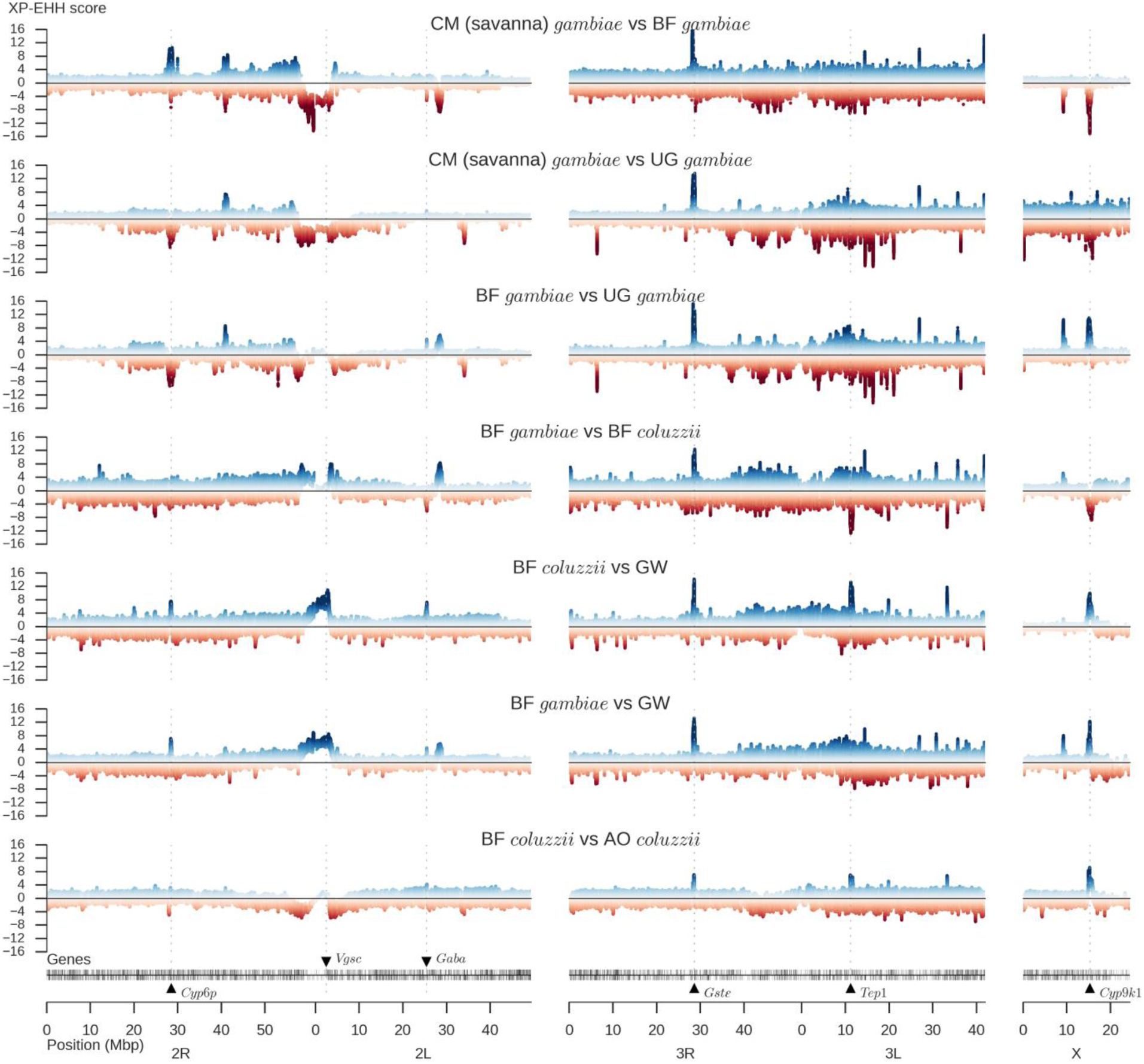
Cross-population genome scans for signatures of recent selection. For each population comparison (e.g., BF gambiae versus BF coluzzii), positive XP-EHH values indicate longer haplotypes and therefore recent selection in the first population (e.g., BF gambiae), and negative XP-EHH values indicate selection in the second population (e.g., BF coluzzii).

**Supplementary Figure 10.**
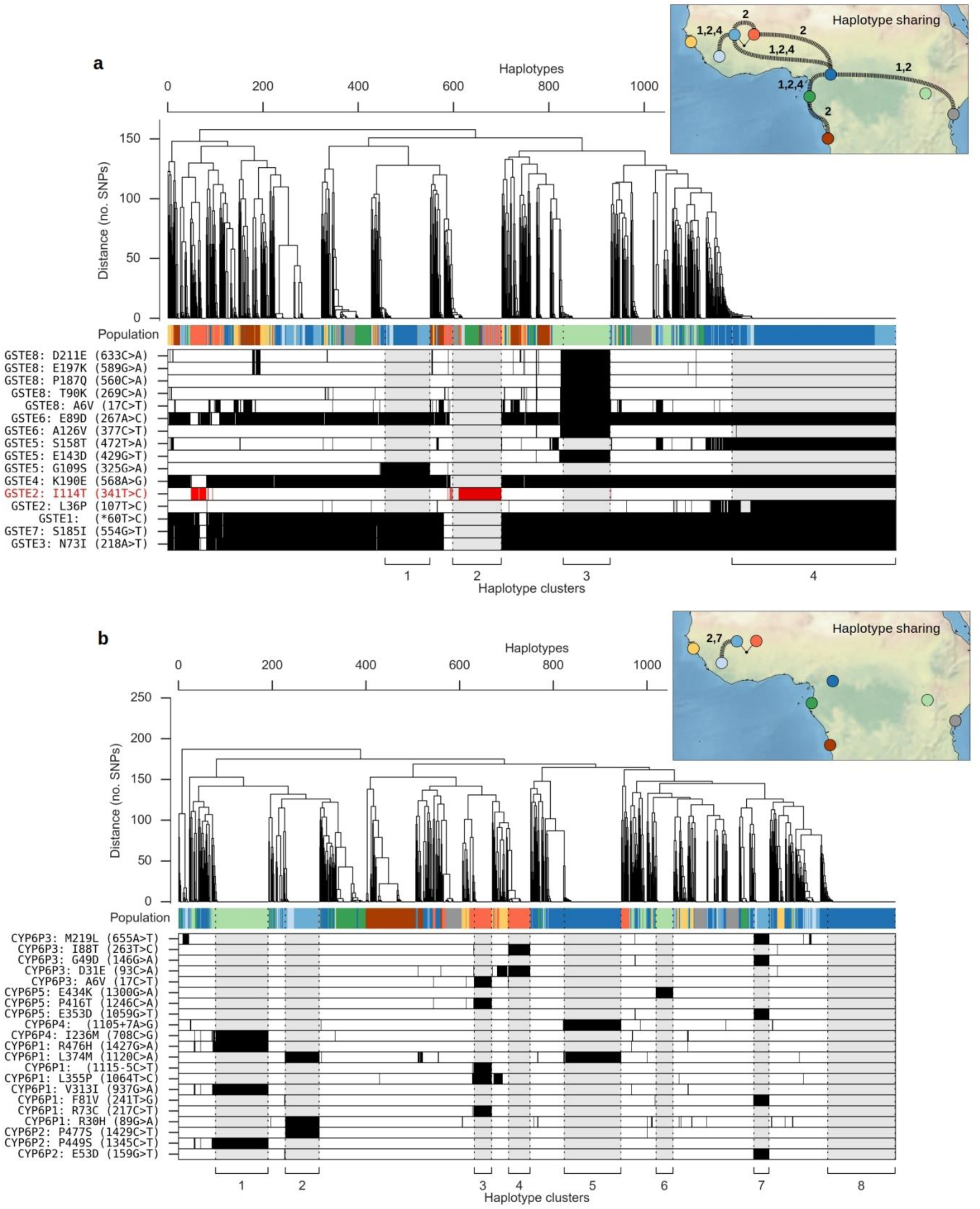
Haplotype structure at metabolic insecticide resistance loci. Plot components are as described for Fig. 5. For both loci, SNPs shown in the lower panel are all either non-synonymous or splice site variants, and are associated with one or more haplotypes under selection. **a**, Haplotype clustering using 1,375 SNPs within the region 3R:28,591,663-28,602,280 spanning 8 genes (Gste1-Gste8). **b**, Haplotype clustering using 1,844 SNPs within the region 2R:28,491,415-28,502,910 spanning 5 genes (Cyp6p1-Cyp6p5).

## Footnotes

‡ http://www.malariagen.net/apps/ag1000g

* http://pubs.usgs.gov/publications/text/East_Africa.html

